# Route of Adenovirus Type 5-Vectored Influenza Vaccination Shapes Systemic and Mucosal Immunity in a Maternal-Neonatal Pig Model

**DOI:** 10.64898/2026.03.12.711389

**Authors:** Stephanie N. Langel, John J. Byrne, Diego Leal, Abigail Williams, Chaitawat Sirisereewan, Danielle Meritet, Michael C. Rahe, Tatiane Terumi Negrão Watanabe, Sasha Compton, Daniela S. Rajao, Juliana Bonin Ferreira, Sean N. Tucker, Elisa Crisci

**Affiliations:** Center for Global Health and Diseases, Department of Pathology, Case Western Reserve University, Cleveland, OH, USA; Department of Population Health and Pathobiology, College of Veterinary Medicine, North Carolina State University, Raleigh, NC, USA; Department of Population Health, College of Veterinary Medicine, University of Georgia, GA, USA; Vaxart, South San Francisco, CA, USA

**Keywords:** pig, influenza virus, maternal immunity, routes of immunization, r-Ad5-HA vaccine

## Abstract

Influenza A virus can cause severe complications in pregnant women and infants, yet no influenza vaccines are approved for infants younger than six months. To address this, novel maternal vaccination strategies are needed to increase global access and coverage in these vulnerable populations. This study evaluated a hemagglutinin (HA) A/California/2009 (H1N1)-based human adenovirus 5 (huAd5) vector vaccine, adjuvanted with a TLR3 agonist, for its ability to induce influenza-specific passive immunity from pregnant and lactating pigs to their piglets following different immunization routes. Influenza naïve pregnant dams were vaccinated via oral, intranasal (IN), or intramuscular (IM) routes three weeks prepartum and boosted four weeks later. Serum, colostrum and milk samples were collected longitudinally to assess HA-specific antibody induced by vaccination. H1N1-Ca/09 neutralizing antibodies were evaluated in serum and IFNγ producing cells were assessed in blood, spleen and lymph node cells. IN and IM routes elicited robust serum HA-specific antibody responses when compared to control animals at one- and four-weeks post-boost, whereas the oral route resulted in poor antibody induction across all samples tested. Piglets nursing from IN and IM vaccinated dams showed a significantly higher level of HA-specific antibodies in serum at 2-3 weeks post-partum compared to control piglets. Notably, IN immunized dams and their piglets showed significantly elevated influenza neutralizing antibodies compared to controls. This work demonstrated that both IN and IM immunization with a huAd5-vectored vaccine robustly induced maternal influenza-specific immunity that supported passive transfer to nursing piglets, with IN immunization resulting in superior transfer of neutralizing antibodies.

## Introduction

Pregnant women and their infants are at increased risk of severe illness caused by influenza viruses (1–4). While immunization of pregnant women with an inactivated trivalent influenza vaccine (IIV) has been recommended by the U.S. Center of Disease Control and Prevention and World Health Organization for over fifteen years, vaccine coverage is suboptimal (5, 6). This challenge is compounded by the fact that IIV immunogenicity is reduced in young children (7, 8), and immunization is not recommended for infants ≤ 6 months, a time when neonates are most vulnerable to influenza-associated mortality (9). Therefore, new and improved influenza vaccine strategies are needed to protect these vulnerable populations. A maternal influenza vaccine that boosts passively transferred influenza-specific antibodies would be an ideal strategy for protection of the mother-neonatal dyad against severe influenza infection.

Maternal antibodies passively transferred across the placenta and into breast milk are critical for protection against infectious disease and immune development during the first year of life (10). For example, protective levels of anti-hemagglutinin (HA) IgG antibodies present in maternal serum are efficiently transferred transplacentally following IIV immunization during pregnancy, and maternal immunization is associated with a reduction in influenza infection among infants (11–13). Breast milk antibodies, particularly secretory IgA, are also associated with decreased episodes of respiratory illness in infants (14). Importantly, most breast milk IgA is derived from IgA⁺ antibody secreting cells primed at distal mucosal sites which subsequently home to the mammary gland during lactation (15, 16). However, because IIV immunization does not stimulate the mucosal immune system directly (17, 18), it is unlikely to promote robust IgA^+^ antibody secreting cell trafficking to the mammary gland and IgA antibody secretion into breast milk. Therefore, we sought to identify vaccination routes capable of stimulating both systemic and mucosal antibody responses in lactating dams, with the goal of maximizing antibody transfer passive transfer and enhancing neonatal protection.

To investigate this strategy, we utilized the pig model, which recapitulates key aspects of human influenza disease, pathogenesis, and immune responses. Pigs are natural hosts of influenza viruses, playing an integral role in the viral ecology, epidemiology and evolution, and they closely resemble humans in anatomy, genetics, physiology, and immune system organization (19, 20). Notably, similar to humans, swine milk contains predominantly IgA making pigs a relevant animal model for the study of maternal-derived milk immunity (10, 21).

We evaluated a replication-defective adenovirus 5 vectored vaccine expressing the influenza HA protein with a novel toll-like receptor 3 agonist as an adjuvant (r-Ad-HA, VXA-A1.1). A previous placebo-controlled, phase 2 clinical trial showed that oral administration of this r-Ad-HA vaccine generated protective immunity in healthy, non-pregnant adults, which was associated with increased circulating IgA^+^ antibody secreting cells and expression of intestinal homing marker β7 on activated plasma cells (22). Additionally, a phase I clinical trial of an Ad5-vectored GI.1/GII.4 norovirus vaccine demonstrated that functional norovirus-specific antibodies were significantly enriched in both breast milk and serum following vaccination during lactation, and that norovirus-specific IgA levels in infant stool increased post-vaccination and positively correlated with breast milk IgA, consistent with passive maternal transfer (23). These findings further support the capacity of mucosal Ad5 platforms to induce functional, maternally transferable antibody responses.

Therefore, the aim of the study was to evaluate the capacity of this r-Ad-HA vaccine to induce systemic and mucosal immunity in pregnant and lactating pigs using different routes of immunization and to assess the passive transfer to their piglets. Influenza naïve pregnant dams were vaccinated via oral, intranasal (IN), or intramuscular (IM) routes three weeks prepartum and boosted four weeks later (one week postpartum). We evaluated HA-specific antibody levels, neutralizing antibody titers and cellular responses in vaccinated dams, in parallel with HA-specific and neutralizing antibodies in their nursing piglets.

Our findings demonstrate that intranasal vaccination of pregnant and lactating dams with the Ad5-vectored influenza vaccine VXA-A1.1 is a highly effective strategy for inducing both systemic and mucosal immunity that is subsequently passively transferred to offspring.

## Materials and methods

### Study design, animals, sample processing

A general experimental layout is summarized in **Figure 1**. A total of sixteen influenza-naive pregnant gilts (Yorkshire × Large White × Landrace cross) were selected for this study. The gilts were sourced and bred at North Carolina State University Swine Educational Unit (SEU), a breeding farm confirmed free of major porcine pathogens, including porcine reproductive and respiratory syndrome virus and influenza A virus.

**Figure 1.**
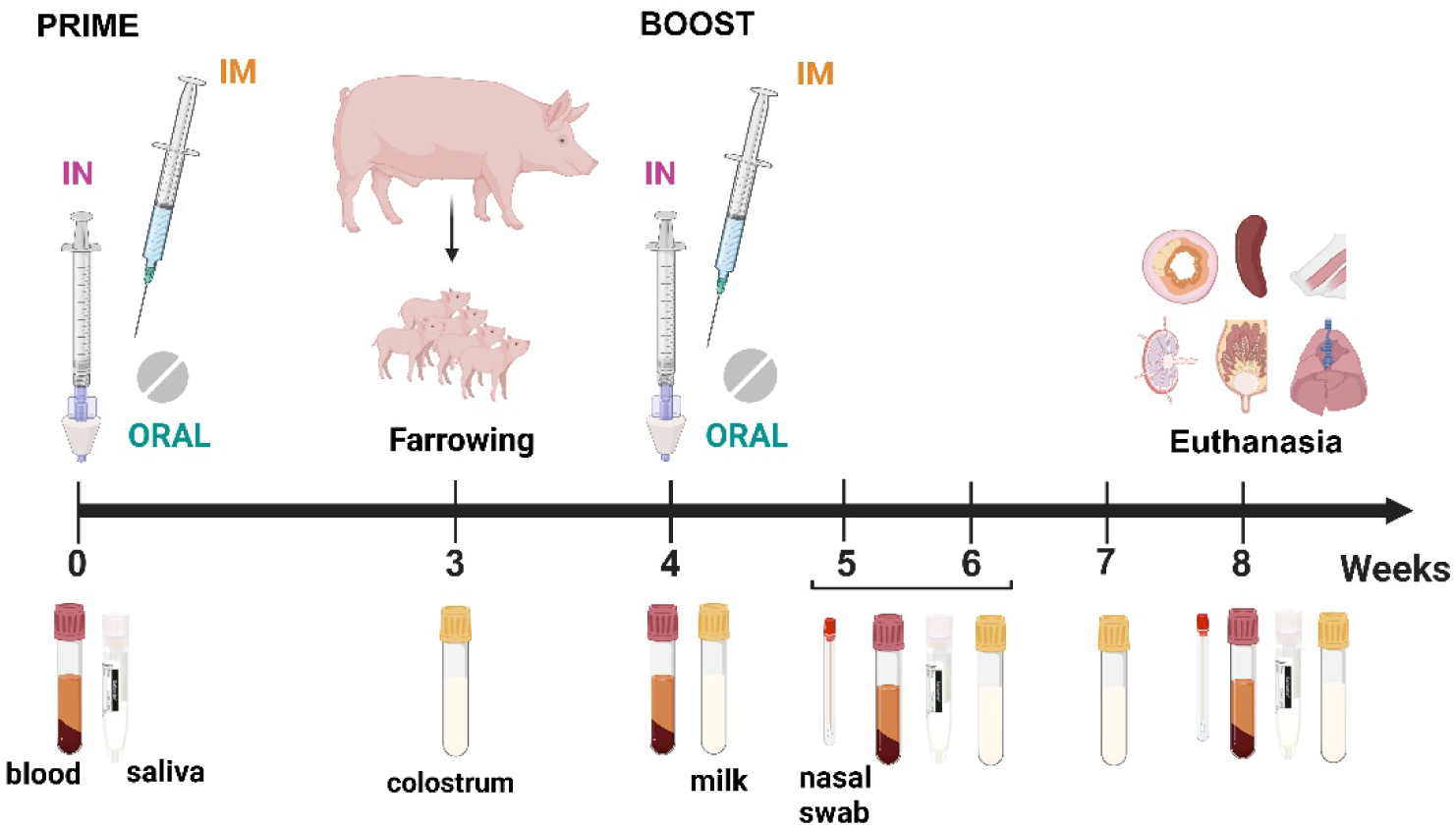
Experimental design and sample collection. Sixteen influenza-naive pregnant gilts (Yorkshire × Large White × Landrace cross) were enrolled and assigned to four groups (n = 4 per group) based on immunization route: intranasal (IN), intramuscular (IM), oral, or unvaccinated control. Dams received a prime immunization 3 weeks prior to farrowing followed by a homologous boost 1 week postpartum (28 days between doses). Hemagglutinin (HA)–specific antibody levels, hemagglutination inhibition (HI) titers, neutralizing antibody titers, and cellular immune responses were assessed in vaccinated dams, together with serum HA-specific antibody levels, HAI titers, and neutralizing antibodies in their nursing piglets. In dams, blood was collected prior to vaccination and at 4, 5–6, and 8 weeks post-prime. In piglets, blood was collected at 2–3 and 5 weeks of age. Saliva, nasal swabs, colostrum, and milk were collected at multiple time points throughout the study. Lungs, mammary gland, lymph nodes, spleen, Peyers’ patches, and nasal mucosa were collected in dams after euthanasia (5 weeks post-partum). Created in BioRender. Crisci, E. (2026) https://BioRender.com/5cbiyk5

Gilts were divided into 4 groups based on the route of immunization used in the study (**Table 1**). Two groups (oral and control) remained at SEU throughout the entire study whereas two groups (IN and IM) were transported to the North Carolina State College of Veterinary Medicine Laboratory Animal Resources (LAR) facilities. Gilts were transported to LAR facilities four weeks before farrowing. All animals tested negative for influenza A virus antibodies (IDEXX Lab, USA) at the beginning and at the end of the study (**Supplementary Figure 1**).

**Table 1.**
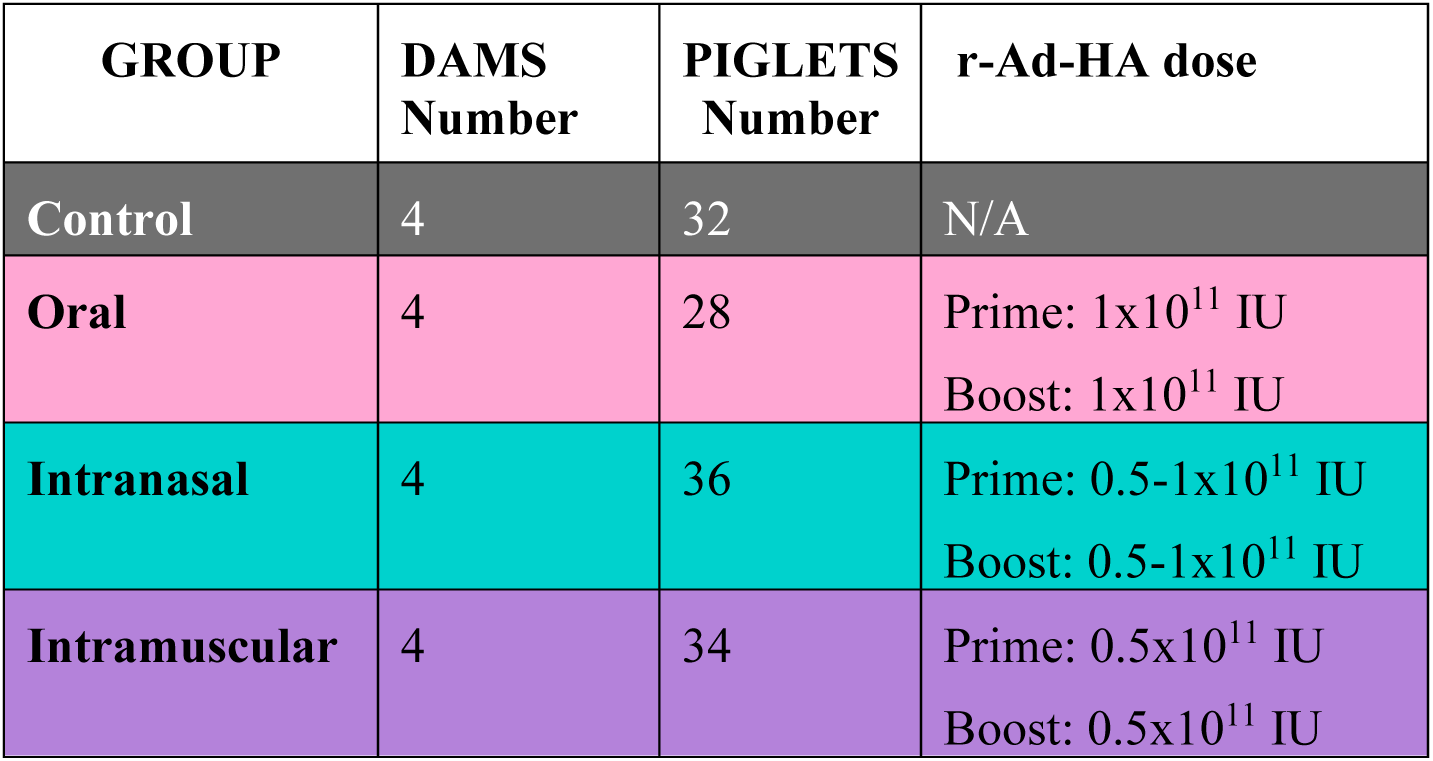
Experimental groups of gilts based on route of immunization.

| GROUP | DAMS Number | PIGLETS Number | r-Ad-HA dose |
| --- | --- | --- | --- |
| Control | 4 | 32 | N/A |
| Oral | 4 | 28 | Prime: $1 \times 10^{11}$ IU<br>Boost: $1 \times 10^{11}$ IU |
| Intranasal | 4 | 36 | Prime: $0.5-1 \times 10^{11}$ IU<br>Boost: $0.5-1 \times 10^{11}$ IU |
| Intramuscular | 4 | 34 | Prime: $0.5 \times 10^{11}$ IU<br>Boost: $0.5 \times 10^{11}$ IU |

Upon arrival at the NCSU CVM LAR facility, dams were housed in individual gestation crates in an environmentally controlled facility. The temperature was maintained at approximately 20°C with a 12-hour light/dark cycle. All gilts had *ad libitum* access to water via a nipple drinker and fed a standard gestation diet twice daily, followed by a lactation diet formulated to meet their nutritional requirements. To ensure a synchronized farrowing schedule, 24-48h before the expected due date, gilts were injected with Lutalyse (Dinoprost 5mg/ml, Zoetis) using the split dose method (1ml/dose, 6h apart, (24)). Farrowing was allowed to proceed naturally.

Within 72h of birth, all piglets underwent standard processing procedures: identification with ear tags or ear notching, iron (IM 200 mg iron dextran, Uniferon® 200 mg/mL, QC) and antibiotic injection (IM, Norocillin 300,000 units/45kg penicillin G as procaine per mL, Norbrook Laboratories or BenzaPen 48 Penicillin G Benzathine and Penicillin G Procaine, Bimeda) in the neck muscle, tail docking and teeth clipping. At SEU piglets also received Spectam Scour Halt oral solution (1 ml, Bimeda). Body weight of each piglet was recorded at birth and on subsequent measurement days.

Dams received a prime immunization dose (IM, IN, oral) 3 weeks before farrowing and a boost dose (IM, IN, oral, respectively) one week after farrowing (**Table 1**). At necropsy, one of the IN inoculated dams was found to be asplenic (25). All data from the asplenic dam was kept in the study. Samples were collected at designated timepoints throughout the study, as summarized in the experimental layout (**Figure 1**). Colostrum was collected during parturition or within 24h after parturition and milk was collected weekly thereafter. Blood collection was performed before each immunization, at 1-2 weeks post boost and at the study endpoint (five weeks postpartum). Nasal swabs were collected using flocked swabs (Copan Diagnostics, USA) and oral fluids were collected with the Salivette® system (Sarstedt, Germany) (**Figure 1**). At necropsy, blood, mammary gland, lymph nodes, lung, nasal mucosa, spleen and Peyer’s patches were collected fresh for mononuclear cell (MNC) isolation or fixed in 10% neutral-buffered formalin solution for histopathological analysis. In piglets, blood was collected at 2-3 weeks after birth and at the study endpoint (five weeks of age).

All experimental procedures involving animals were conducted in strict accordance with the guidelines established by the Institutional Animal Care and Use Committee (IACUC) of North Carolina State University and were approved under protocols #22-122 and #20-473.

### Vaccine

The vaccine consisted of a replication-deficient, E1/E3-deleted adenovirus type 5 (Ad5) vector expressing the HA protein from influenza A/California/04/2009 (H1N1), similar to that previously described (22). The vaccine was produced under non-GMP conditions for use in animal studies.

### Virus and cells

Influenza A/California/04/2009 H1N1 virus (BEI Resources, NR-13659) was propagated in MDCK-SIAT1 cells (CRL-3742, ATCC). Viral stocks were submitted for sequencing for confirmation of strain by an outsourced sequencing service (Cambridge Technologies). This virus was then used in all immunological assays. Viral titers were determined using the immunoperoxidase monolayer assay (IPMA) and expressed as TCID50/mL as determined by the Reed and Muench method (26).

### Immunoperoxidase monolayer assay (IPMA)

Influenza A virus infected MDCK-SIAT1 cells were fixed with a 80% acetone solution for 15 minutes at room temperature, then washed twice with phosphate-buffered saline containing 0.1% Tween-20 (0.1% PBST) and subsequently blocked with 3% bovine serum albumin in PBS for 1 hour at 37°C. Influenza A virus was detected by a mouse anti-H1N1-NP-protein IgG1 monoclonal antibody (clone IC5-1B7, BEI Resources, NR-43899) (1:1000) after incubation for 1 hour at 37°C. After two washes with 0.1% PBST, the cells were incubated with HRP-conjugated rabbit anti-mouse antibody (ab6728, Abcam, USA), diluted 1:3000, for 1 hour at 37°C. Following two additional washes with 0.1% PBST, the cells were counterstained with 3-amino-9-ethylcarbazole (AEC) substrate. Following substrate development, infected cells were visualized by light microscopy, and viral titers were determined by identifying the highest dilution yielding detectable antigen-positive cells, expressed as the tissue culture infectious dose (TCID₅₀).

### ELISA

Antibody detection was performed via an in-house HA-specific ELISA. Flat bottom 96-well Microtiter™ Microplates (Thermo scientific) were coated with 50µL per well of a 1 µg/mL stock of purified A/California/04/2009 HA-trimer (Sino Biological) in 1x KPL Coating Solution (Fisher) and incubated at 4°C overnight. Plates were washed 5 times with 0.1% PBST after each incubation. Protein coated plates were blocked with 50µL blocking buffer (1% fish gelatin, 1% normal goat serum, 2% bovine serum albumin in 0.1% PBST) and incubated at 37°C for 1 hour. To evaluate immunoglobulins in serum, milk and colostrum, samples were diluted 1:300 in blocking buffer, added to the plate at 50µL per well, and incubated at 37°C for 1 hour. To evaluate immunoglobulins in oral fluid and nasal swabs, samples were diluted 1:10 in blocking buffer, added to the plate at 50µL per well, and incubated at 37°C for 1 hour. To evaluate immunoglobulins in piglet serum, samples were diluted 1:100 in blocking buffer, added to the plate at 50µL per well, and incubated at 37°C for 1 hour. Antibody binding was detected by adding 50µL per well of goat anti-porcine IgG (Abcam) or IgA (Bethyl) HRP-conjugated antibodies diluted 1∶4000 in blocking buffer at 37°C for 1 hour. The colorimetric detection was evaluated using 80µL per well of KPL SureBlue Reserve™ TMB Microwell Substrate (VWR) per well. Following a 15-minute incubation at room temperature, the reaction was stopped by addition of 80µL per well of KPL SureBlue Reserve™ STOP (VWR). Optical density was measured at 450 nm using a Biotek Microplate spectrophotometer.

The IDEXX Swine Influenza Virus Antibody Test ELISA (nucleoprotein [NP] antigen) was performed according to the manufacturer’s specifications.

### Virus neutralization assay

Serum neutralizing antibody detection was performed in flat bottom 96-well tissue culture plates (Genesee Scientific). Samples were prediluted 1:100 in DMEM (Genesee Scientific) without additives and plated at 55.5µl/well. All samples were further diluted directly in the plate using six 10-fold serial dilutions in DMEM without additives. Wells containing DMEM alone (no sample) were included as negative controls. All wells were adjusted to a final volume of 50µL. H1N1 Ca/09, previously propagated in MDCK-SIAT1 cells, was diluted to a concentration of 200 TCID_50_/50µL. The plate was incubated at 37°C, with 5% CO_2_, for 2 hours. MDCK-SIAT1 cells were then added to the plate at a concentration of cells/100µL in each well and incubated for 18 hours at 37°C, with 5% CO_2_. The plates were then fixed and analyzed utilizing the IPMA protocol (section 2.3). Neutralizing antibodies titers were recorded as the highest serum dilution yielding no detectable antigen-positive cells.

### Hemagglutination inhibition (HI) assay

Serum samples were analyzed by HI assay using standard techniques (27). Briefly, sera were treated with Receptor Destroying Enzyme (RDE; Denka Seiken, Tokyo, Japan) overnight at 37°C followed by inactivation at 56°C for 30 min and adsorption with 0.5% turkey red blood cells (RBCs) for 30 min. Final sample dilution was 1:10 in 0.85% Saline solution. Two-fold serially diluted sera were mixed in equal volumes (25uL each) with 4 hemagglutination units (HAU) of H1N1 Ca/09 virus. The serum–virus mixtures were incubated at room temperature for 30 minutes, after which 0.5% turkey red blood cells (RBCs) were added and incubated for an additional 30 minutes. HI titers were recorded as the highest serum dilution that completely inhibited hemagglutination.

### IFNγ ELISpot assay

The porcine interferon gamma (IFNγ) ELISpot assay was performed as previously described, with minor modification (28). 96-well filter plates with hydrophobic polyvinylidene fluoride membranes (Millipore, USA) were treated with 50µL 70% ethanol for 1 minute. After activation, the plates were washed five times with ddH2O and coated 100µL per well overnight at 4°C with a monoclonal antibody specific to porcine IFNγ (1:50; Mabtech, Nacka Strand, Sweden). The antibody-coated plates were washed with PBS and incubated with stimulants, including the positive control (Concanavalin A, 5 µg/mL), and influenza strain A/California/04/2009 at an MOI of 0.5. The wells containing only cells and complete media served as mock controls. All cells were thawed, washed with RPMI-1640 medium, counted using trypan blue, and seeded at 250,000 cells/well. The plates were then incubated at 37°C in 5% CO_2_ overnight, then washed with PBS and incubated with a biotinylated IFNγ-specific antibody (clone P2C11, Mabtech, Nacka Strand, Sweden) at 1 µg/mL for 1 hour at room temperature. After washing, streptavidin-alkaline phosphatase (Mabtech, Sweden) was added at a 1:2000 dilution and incubated for 60 minutes at room temperature. Then 100 µl alkaline phosphatase substrate, 5-bromo-4-chloro-3-indolyl phosphate/nitro blue tetrazolium (BCIP/NBT; Sigma-Aldrich, Germany) was added to the plate and incubated for 25 minutes. The reaction was stopped by rinsing with DI H2O water, and the plates were left to dry overnight. Spot counting was performed using the Mabtech ELISpot reader (Mabtech, NackaStrand, Sweden). IAV-specific IFNγ producing cells were expressed as spot-forming units (SFU) per million cells.

### CD3 and CD20 Immunohistochemistry and grading analysis

Immunohistochemistry staining was performed on the Biocare Intellipath FLX by the histology core at the North Carolina State University College of Veterinary Medicine. Briefly, 4 μm tissue sections on positively charged slides were baked for 30 minutes at 60℃. Slides were then deparaffinized followed by antigen retrieval in citrate buffer (pH 6.0). Endogenous peroxidase activity was blocked with 3% hydrogen peroxide for 20 minutes at room temperature and then washed with a TBS buffer rinse. Fc blocker (Innovex Biosciences) was added for 30 minutes at room temperature, followed by a TBS buffer rinse. The primary antibody for CD3 staining was a rabbit polyclonal (Dako A0452) used at a dilution of 1:200 for 30 minutes of incubation at room temperature. This was followed by a TBS buffer rinse. For CD20 staining a rabbit polyclonal (Invitrogen PA5-16701) was used at a dilution of 1:600 for 30 minutes of incubation time at room temperature, with subsequent TBS buffer wash. For both primary antibodies, a polymer rabbit HRP (Vector Labs) was used as the secondary antibody with an incubation time of 45 minutes for CD3 staining and 35 minutes for CD20 staining at room temperature, followed by a TBS buffer rinse. DAB chromogen (Vector labs) was incubated on the slide for 4 minutes and rinsed with dH2O, followed by hematoxylin stain for 5 minutes and one final TBS buffer wash. Slides were dehydrated through alcohol and xylene, then coverslipped.

Grading of lymphocytes was performed by a board certified veterinary anatomic pathologist. Lymphocyte counts (T, B, or plasma cells) were averaged across 10 high-power fields (hpf) per tissue type. Grade 0 = No T/B cells or plasma cells present in any hpf, Grade 1 = <10 cells per hpf, Grade 2 = 10-50 cells per hpf, Grade 3 = >50 cells per hpf.

### Statistical analysis

All analyses were performed using GraphPad Prism version 10.5.0 for Windows (GraphPad Software Inc., San Diego, CA, USA). Data are presented as mean ± SEM, and a p value < 0.05 was considered statistically significant. Longitudinal antibody responses in dams were analyzed using two-way ANOVA with Bonferroni’s multiple-comparison test. Comparisons among vaccination groups for piglet antibody responses, neutralizing and HI titers, and cellular immune responses were performed using non-parametric Kruskal–Wallis tests followed by Dunn’s multiple-comparison correction. Correlations between maternal and piglet antibody levels were assessed using simple linear regression analysis, with goodness-of-fit reported as the coefficient of determination (R²). All tests were two-tailed.

## Results

### IN maternal r-Ad-HA vaccination induced robust systemic IgG responses

Sows were vaccinated with r-Ad-HA using a prime-boost regimen, with the prime dose administered three weeks prepartum and the booster dose given one week postpartum. Serum HA-specific IgG levels were significantly elevated in the IN group compared to the control and oral groups at one- and four-weeks post-boost (**Figure 2A**). Although the IM group showed elevated serum HA-specific IgG, one markedly low-responder animal increased variance sufficiently to prevent the group from reaching significance (**Figure 2A**). In contrast, serum HA-specific IgA levels were not significantly different between groups at any time point (**Figure 2B**).

**Figure 2.**
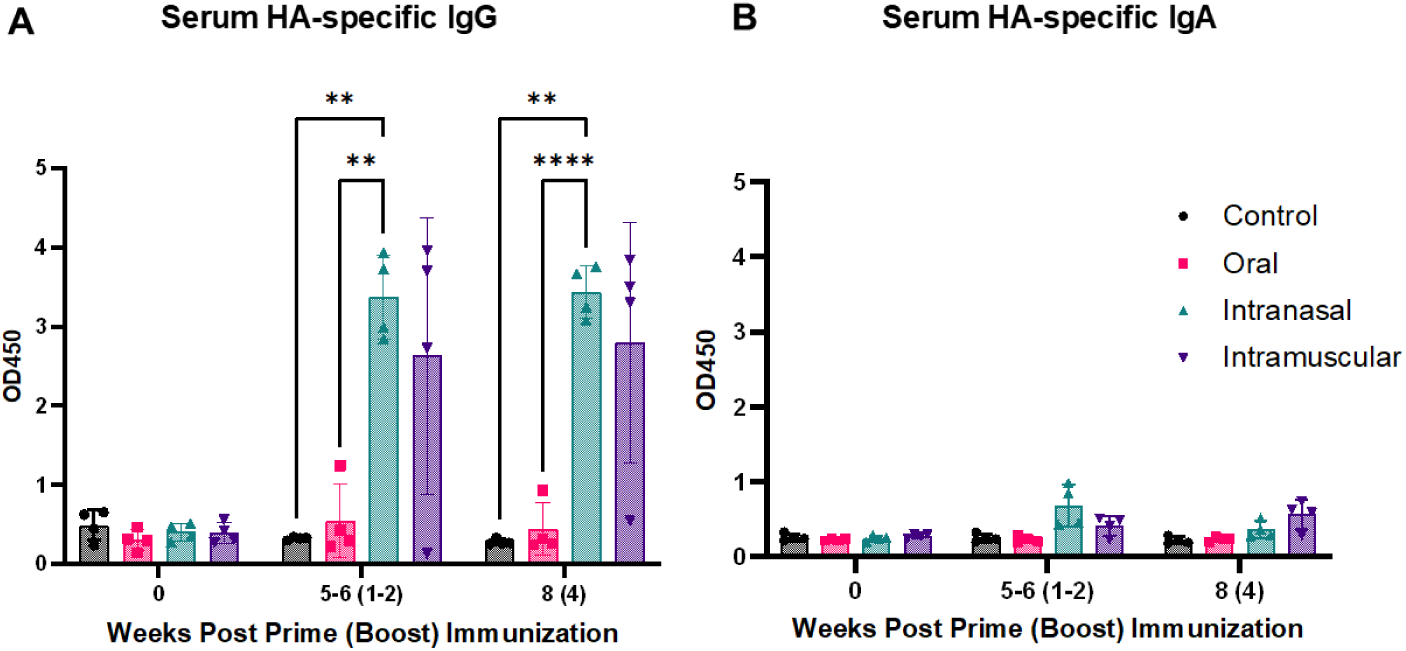
Maternal vaccination with r-Ad-HA induces a significant increase in serum HA-specific IgG following both intranasal (IN) and intramuscular (IM) administration. Dam serum HA-specific IgG (A) and IgA (B) antibody levels were determined by ELISA. Data are shown as mean ± SEM. The x-axis indicates weeks post-prime immunization, with weeks post-boost administration indicated in parentheses (e.g., 8 (4)). Significance was determined by a two-way ANOVA with Bonferroni’s test for multiple comparisons (**p < 0.01, ****p < 0.0001).

### IN and IM maternal r-Ad-HA vaccination increased colostral IgG, while only IM delivery increased IgG and IgA in early mature milk

Colostrum HA-specific IgG levels were significantly elevated in both the IN and IM groups compared to control animals (**Figure 3A**). In contrast, only one of four orally vaccinated dams exhibited elevated HA-specific IgG in colostrum (**Figure 3A**). Colostrum HA-specific IgA levels were elevated in both IN and IM groups; however, neither showed a statistically significant increase compared to the control group (**Figure 3B**), likely due to elevated background levels in a single control dam (**Figure 3B**). Milk HA-specific IgG and IgA levels were significantly increased in the IM group, but only during early lactation at 4 weeks post-prime (day of boost) (**Figure 3A, B**). By 6 weeks post-prime (2 weeks post-boost), HA-specific IgG levels demonstrated a trend toward difference in the IM group compared to control and oral groups (p = 0.05–0.1).

**Figure 3.**
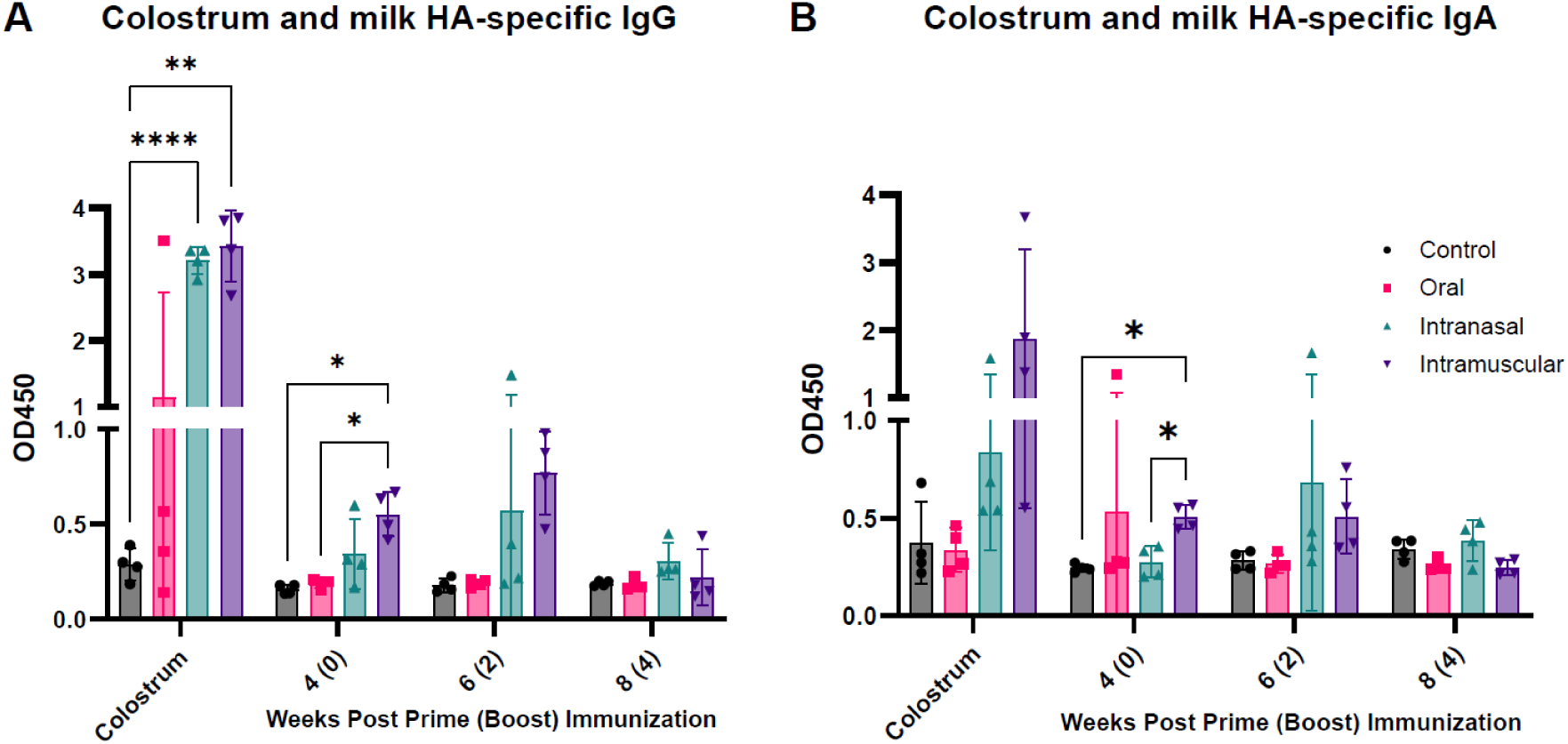
Maternal r-Ad-HA vaccination significantly increases colostrum HA-specific IgG levels following both intranasal (IN) and intramuscular (IM) administration, while only IM delivery increases IgG and IgA in early mature milk. Dam colostrum and milk HA-specific IgG (A) and IgA (B) antibody levels were determined by an ELISA. Data shown as mean ± SEM. The x-axis indicates weeks post-prime immunization, with weeks post-boost administration indicated in parentheses (e.g., 8 (4)). Significance was determined by a two-way ANOVA with Bonferroni’s test for multiple comparisons (*p < 0.05, **p < 0.01, ****p < 0.0001).

### HA-specific IgG and IgA in nasal and oral secretions were not increased by any vaccination route

Although IN immunization modestly increased HA-specific IgA levels in nasal secretions, neither HA-specific IgG nor IgA reached significant levels in nasal or oral secretions compared to control animals (**Supplementary Figure 2**).

### IN and IM maternal r-Ad-HA vaccination conferred passive antibody immunity to suckling piglets

At 1-2 weeks post-boost, piglets born to IN- and IM-vaccinated dams had significantly higher HA-specific IgG levels compared to those born to control or orally vaccinated dams, with IM piglets also exhibiting significantly higher levels than IN piglets. (**Figure 4A**). Serum HA-specific IgA levels were significantly higher in IM piglets than in all other groups, with IN piglets also exhibiting significantly higher levels than the oral group. (**Figure 4B**). Dam colostrum and early milk HA-specific IgG concentrations were positively correlated with piglet serum IgG levels, with the strongest association observed at week 4 (R² = 0.81, p < 0.0001), followed by a progressive decline in association over time (**Table 2**). In contrast, HA-specific IgA correlations were largely restricted to colostrum (R² = 0.66, p < 0.0001), with milk IgA showing no significant association at weeks 4 or 6 and a weak inverse association emerging at week 8 (**Table 2**).

**Figure 4:**
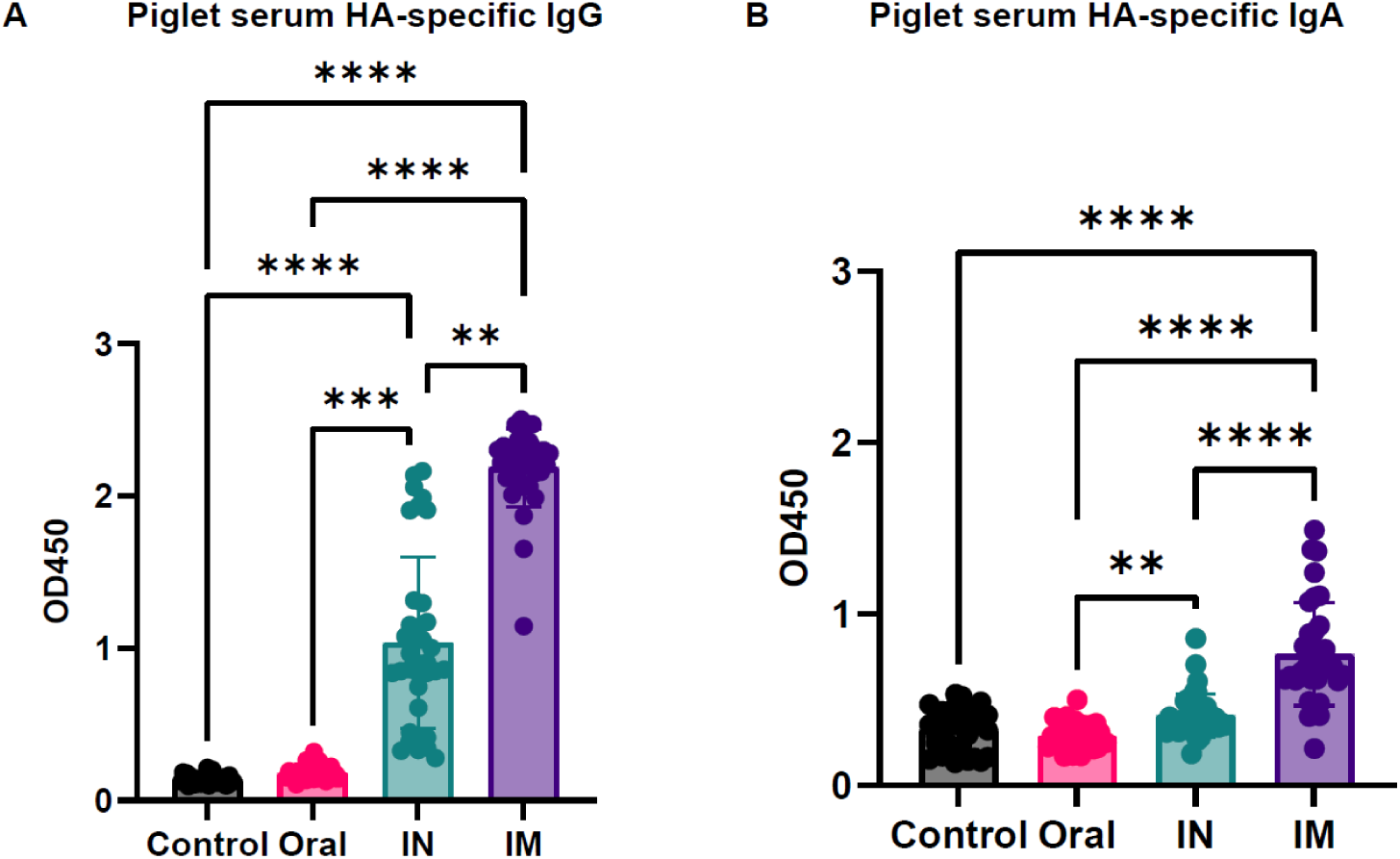
IN and IM maternal r-Ad-HA vaccination conferred passive antibody immunity to suckling piglets. Serum HA-specific IgG (A) and IgA (B) antibody levels in piglets were determined by ELISA. Data are shown as mean ± SEM. Statistical significance was determined using a Kruskal–Wallis test with Dunn’s multiple-comparison correction (**p < 0.01, ***p < 0.001, ****p < 0.0001).

**Table 2.** Correlations between sow milk HA-specific IgG/IgA and piglet serum HA-specific IgG/IgA.

| Isotype | Maternal Compartment | Direction | R<sup>2</sup> | P value |
| --- | --- | --- | --- | --- |
| IgG | Colostrum | Positive | 0.532 | <0.0001 |
|  | Milk Week 4 | Positive | 0.815 | <0.0001 |
|  | Milk Week 6 | Positive | 0.331 | <0.0001 |
|  | Milk Week 8 | Negative | 0.089 | 0.0006 |
| IgA | Colostrum | Positive | 0.656 | <0.0001 |
|  | Milk Week 4 | Positive | 0.001 | 0.7222 |
|  | Milk Week 6 | Positive | 0.014 | 0.1787 |
|  | Milk Week 8 | Negative | 0.032 | 0.0407 |

### IN maternal vaccination boosts dam virus neutralization and HI titers

At 1–2 weeks post-boost, serum neutralizing titers in IN-vaccinated dams were significantly higher than those in the control and oral groups (**Figure 5A**). Piglets born to IN- and IM-vaccinated dams also exhibited significantly higher neutralizing titers compared to control and oral groups (**Figure 5B**), with piglets from IN-vaccinated dams showing the highest responses, exceeding those of IM piglets (**Figure 5B**). HI assays demonstrated that IM- and IN-vaccinated dams had higher HI titers than control and orally vaccinated animals, but differences were not significant (**Figure 5C**). Although piglets nursing from IN-vaccinated dams displayed higher HI titers, these differences did not reach statistical significance compared to other groups (**Figure 5D**). Dam and piglet neutralizing antibody titers were positively correlated; however, the strength of the association was modest (R² = 0.19), indicating considerable variability between maternal and neonatal responses.

**Figure 5.**
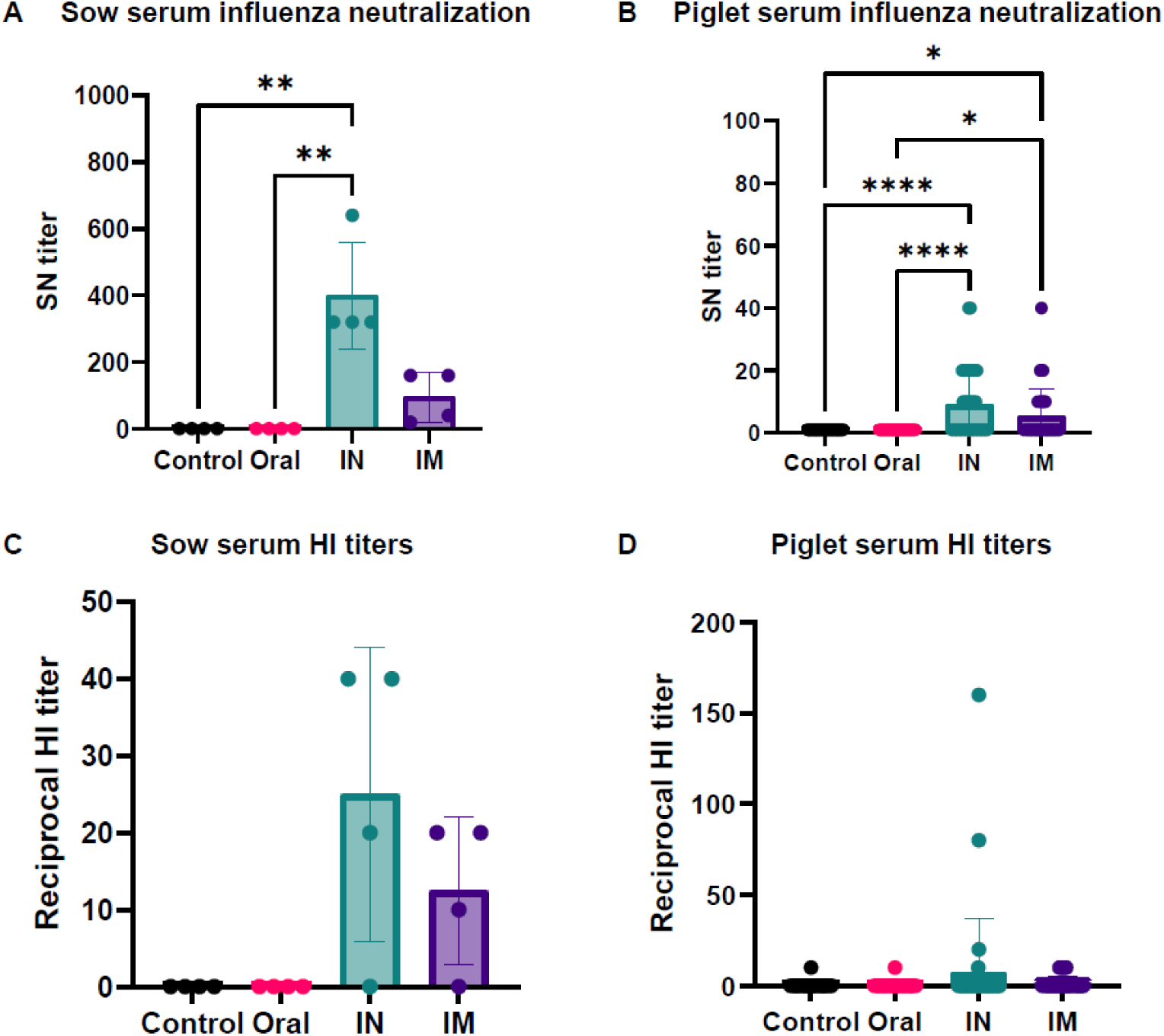
Maternal vaccination route shapes neutralizing and hemagglutination inhibition antibody responses in dams and their piglets. (A) Dam and (B) piglet serum influenza neutralization titers were measured using a microneutralization assay. (C) Dam and (D) piglet hemagglutination inhibition (HI) titers were measured in parallel. Data are presented as mean ± SEM. Statistical significance was determined using a one-way Kruskal–Wallis test followed by Dunn’s multiple-comparisons test. (*p < 0.05, **p < 0.01, ****p < 0.0001).

### Cellular immunity

Systemic and tissue-specific cellular immunity was evaluated in dams at the study endpoint, 4 weeks post-vaccination boost (**Figure 6**). For systemic responses, IN immunized dams showed significantly higher IFNγ-producing peripheral blood mononuclear cells (PBMCs) compared with IM-immunized and control animals (**Figure 6A**). In the spleen, IM-vaccinated animals exhibited significantly higher numbers of IFNγ-producing cells compared with control and orally vaccinated dams, a difference driven by a single IM-vaccinated outlier (**Figure 6B**). In the submandibular lymph node, cellular responses did not differ significantly between groups (**Figure 6C**).

**Figure 6:**
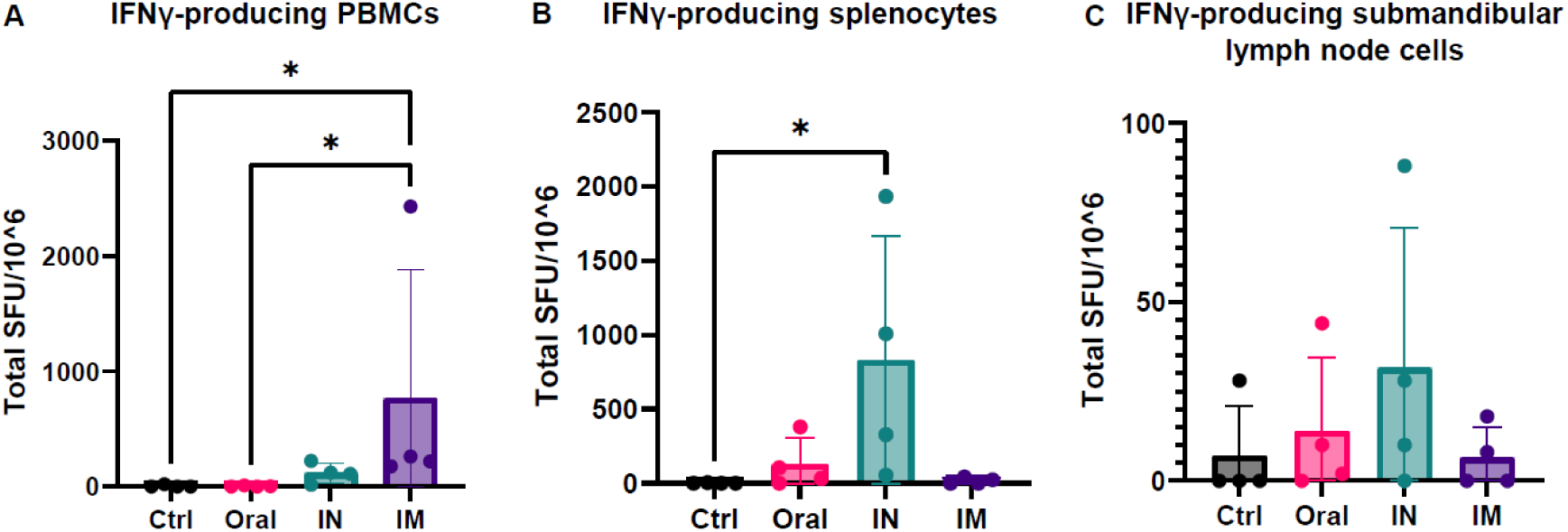
Systemic and tissue-specific cellular immunity in vaccinated dams. Systemic and tissue-specific cellular responses were assessed at the study endpoint, 4 weeks post–vaccination boost. (A) IFNγ-producing PBMCs, (B) IFNγ-producing splenocytes, and (C) IFNγ-producing submandibular lymph node cells were quantified by ELISpot. Data are presented as mean ± SEM. Statistical significance was determined using a Kruskal–Wallis test with Dunn’s multiple-comparison correction (*p < 0.05).

### Immunohistochemistry data

Lymphocytic infiltration (T cells, B cells, and plasma cells) of the lung and mammary gland were evaluated for all pigs at necropsy. For the evaluation of T and B cells, CD3 and CD20 immunohistochemistries were performed. Plasma cells can be distinguished from other lymphocytes and enumerated on H&E-stained slides. There were no statistical differences in infiltrating lymphocyte types in any of the evaluated tissues based on route of vaccination (**Supplemental figure 3**).

## Discussion

Vaccine strategies that improve maternal protection while simultaneously increasing passive immunity in the infant are critical for safeguarding these vulnerable populations (29). Enhancing transplacental transfer of IgG requires elevated maternal serum IgG levels, whereas increasing antibody levels in breast milk, particularly IgA, depends on stimulating robust mucosal immune responses (30). To determine the optimal vaccination route for inducing both systemic and mucosal immune responses, we evaluated a r-Ad5-HA vaccine administered to pregnant and lactating pigs via three routes: oral delivery as an enteric-coated pill, and IN or IM administration as liquid formulations. Adenovirus serotype 5 vectors require engagement of the host coxsackie and adenovirus receptor (CAR) to deliver the vaccine transgene. In this construct, the transgene encodes influenza HA, which is translated into infected host cells and subsequently processed for immune recognition. The r-Ad5 vector is replication-deficient due to deletions in the E1 and E3 regions, thereby preventing the generation of progeny virions. These characteristics have established Ad5 vectors as a widely used and immunogenic vaccine platform across multiple infectious disease applications (31, 32). Previous, but a limited number of studies, have demonstrated that Ad5 vectored vaccine platforms can boost maternal immunity against RSV in cotton rats (33), porcine epidemic diarrhea virus (PEDV) in pigs (34), cytomegalovirus in guinea pigs (35), and Zika virus in mice (36). Using this platform, we employed a maternal–neonatal pig model to determine whether maternal influenza immunization enhanced route-dependent passive antibody transfer to suckling piglets.

Both IN and IM immunization induced robust HA-specific serum IgG responses; however, only IN immunization resulted in significantly higher IgG levels compared with control and orally vaccinated dams at both 1–2 and 4 weeks post-boost. Similar observations have been reported in humans. For example, in heterologous booster studies of adults previously primed with an inactivated COVID-19 vaccine, both aerosolized IN and IM Ad5-nCoV boosters elicited comparable increases in receptor-binding domain–specific IgG levels, with no significant differences between routes of administration, highlighting the capacity of mucosal boosting to reinforce systemic IgG responses (37, 38). Interestingly, vaccination of pregnant dams at 5 and 2 weeks prepartum with an r-huAd5 vector expressing the spike protein of PEDV resulted in significantly higher spike-specific IgG responses following IM/IM immunization compared to IN/IN administration (34). The differences observed between IM and IN Ad5 vaccination routes in our study compared to theirs may reflect antigen-specific effects, differences in immunization regime or differences in host genetics, as our dams were bred in the United States whereas theirs were bred in China. Interestingly, we observed no increase in HA-specific IgG above control levels in three of four orally immunized dams. This may reflect limited CAR expression in the porcine intestinal epithelium, leading to reduced or incomplete adenoviral transgene delivery (39).

Serum HA-specific IgA levels were only modestly increased in some animals at the time points measured, although not statistically. This may reflect the fact that antigen-specific IgA induced by IN is predominantly produced by tissue-resident plasma cells and secreted locally as secretory IgA via polymeric Ig receptor–mediated transport (40), resulting in limited contribution to circulating IgA pools (41).

We next assessed HA-specific IgG and IgA levels in the colostrum and milk of immunized dams. Colostrum, produced during the first 24 to 48 hours after birth, is thicker and richer in fat compared to mature milk and contains higher concentrations of antibodies and immune cells (10). Unlike humans, pigs have an epitheliochorial placenta that does not permit transplacental antibody transfer. As a result, piglets rely entirely on colostrum for the acquisition of maternal antibodies into circulation (42). We observed significantly higher levels of HA-specific IgG in the colostrum of IN and IM vaccinated dams compared with control and orally immunized animals. In pigs, colostral IgG is derived almost entirely from maternal serum via selective transport into the mammary gland (43). Accordingly, the vaccination routes that elicited robust HA-specific IgG responses in the circulation (IN and IM) also resulted in significantly elevated HA-specific IgG levels in colostrum and early mature milk. Notably, only one of the four orally immunized dams exhibited HA-specific IgG levels comparable to those observed in the IN and IM groups, suggesting incomplete effectiveness of the oral immunization strategy. Despite the presence of a single high-binding control animal that increased variability, HA-specific IgA levels in colostrum followed a similar trend to IgG, with IN- and IM-vaccinated dams demonstrating higher responses compared to oral and control groups, although these differences did not reach statistical significance. Early studies suggest that approximately 40% of IgA in pig colostrum is derived from serum (43). Therefore, the elevated HA-specific IgA in the IN group may represent the combined contribution of locally produced mucosal IgA and serum transudation, whereas IgA in the IM group would primarily reflect systemic responses alone.

In piglet serum, HA-specific IgG and IgA levels were significantly higher in offspring born to IM-immunized dams compared to all other groups. Although colostrum HA-specific IgG levels were overall comparable between IN- and IM-vaccinated dams, three of the four IM-vaccinated dams exhibited numerically higher IgG levels than those in the IN group. Consistent with the colostrum-dependent transfer of maternal IgG in pigs, piglet serum IgG levels mirrored this pattern. Similarly, although HA-specific IgA was present at lower concentrations than IgG in both compartments, piglet serum IgA levels paralleled the trend observed in dam colostrum, with IM-immunized dams exhibiting numerically higher responses than the IN group. Additionally, each dam had an average litter size of approximately 8 piglets, resulting in group sizes ranging from 28 to 39 piglets. This larger sample size, relative to the dam dataset, increased statistical power in the piglet analyses and improved sensitivity to detect differences between treatment groups.

Although piglets born to IM-immunized dams had significantly higher levels of HA-specific IgG and IgA in serum, those born to IN-immunized dams exhibited significantly higher influenza neutralizing antibody titers, showing a difference in antibody functionality. This pattern was mirrored in maternal serum at one week postpartum, shortly after the transition from colostrum to mature milk, where only IN-immunized dams showed significantly elevated neutralizing titers compared to control and orally immunized groups. This likely reflects greater functional neutralizing capacity against the tested strain in IN-immunized dams and piglets, potentially due to qualitative differences such as more effective epitope targeting or higher affinity. Supporting this, HAI titers, which reflect the ability of antibodies to block viral attachment to sialic acid receptors on red blood cells, were significantly higher in IN-immunized dams compared to oral and control groups. However, no significant differences in HAI titers were observed among piglets, consistent with the lower sensitivity of the HAI assay relative to the virus neutralization assay observed in other studies (44, 45). These data suggest that, when administered using the same vaccine platform, IN vaccination may be more effective than IM vaccination at eliciting antibodies with enhanced functional anti-influenza activity. This may be due to multiple factors, including route-dependent differences in antigen expression and tissue tropism, distinct innate immune programming and germinal center dynamics, potential differences in isotype distribution, variations in epitope targeting and affinity maturation, and qualitative features such as antibody glycosylation or avidity that collectively shape functional neutralization capacity. These possibilities warrant further mechanistic investigation to better define how vaccination route shapes antibody quality.

We next assessed IFN-γ production by MNCs isolated from PBMCs, spleen, and submandibular lymph nodes as a measure of T cell activation. IN-immunized dams exhibited significantly higher IFN-γ production from PBMCs compared to IM-immunized dams, whereas the opposite was observed in splenic mononuclear cells, where IFN-γ production was greater in the IM group. This compartmentalized pattern may reflect differences in T cell trafficking induced by the route of immunization. IN vaccination likely promotes the generation of mucosa-homing T cells that circulate through the blood, contributing to elevated IFN-γ responses in PBMCs. In contrast, IM immunization may result in greater antigen drainage to systemic lymphoid tissues such as spleen, leading to enhanced splenic T cell activation. IFN-γ responses in the submandibular lymph nodes more closely mirrored those observed in the blood, with the highest levels detected after IN immunization, although the differences were not statistically significant. This likely reflects the anatomical location of these lymph nodes at the base of the jaw, where they drain lymph from the upper respiratory tract and oral cavity, positioning them to filter and respond to antigens delivered via the IN route.

## Summary and conclusions

This study demonstrates that the route of immunization critically shapes the functional properties of milk-transferred antibodies. Using a maternal–neonatal pig model, we show that IN immunization outperforms IM delivery in generating antibodies with enhanced functional activity in the neonatal circulation. Moreover, the route of immunization influenced the anatomical distribution of T cell responses, with IN delivery favoring mucosal and circulating compartments, whereas IM delivery enhanced responses in the spleen. Notably, differential routes of Ad5-based influenza vaccination have not previously been evaluated in a maternal–neonatal system, highlighting the novelty of this work. Together, these findings underscore the importance of vaccine delivery route in shaping maternal immune responses and support IN vaccination as a promising complementary strategy to improve maternal influenza immunization and neonatal protection. Maternal antibody transfer through breast milk may help overcome key limitations of direct newborn immunization by leveraging the mother’s mature immune system to provide protection during the period when infants are most vulnerable to infection.

---

**Supplementary Figure 1:**
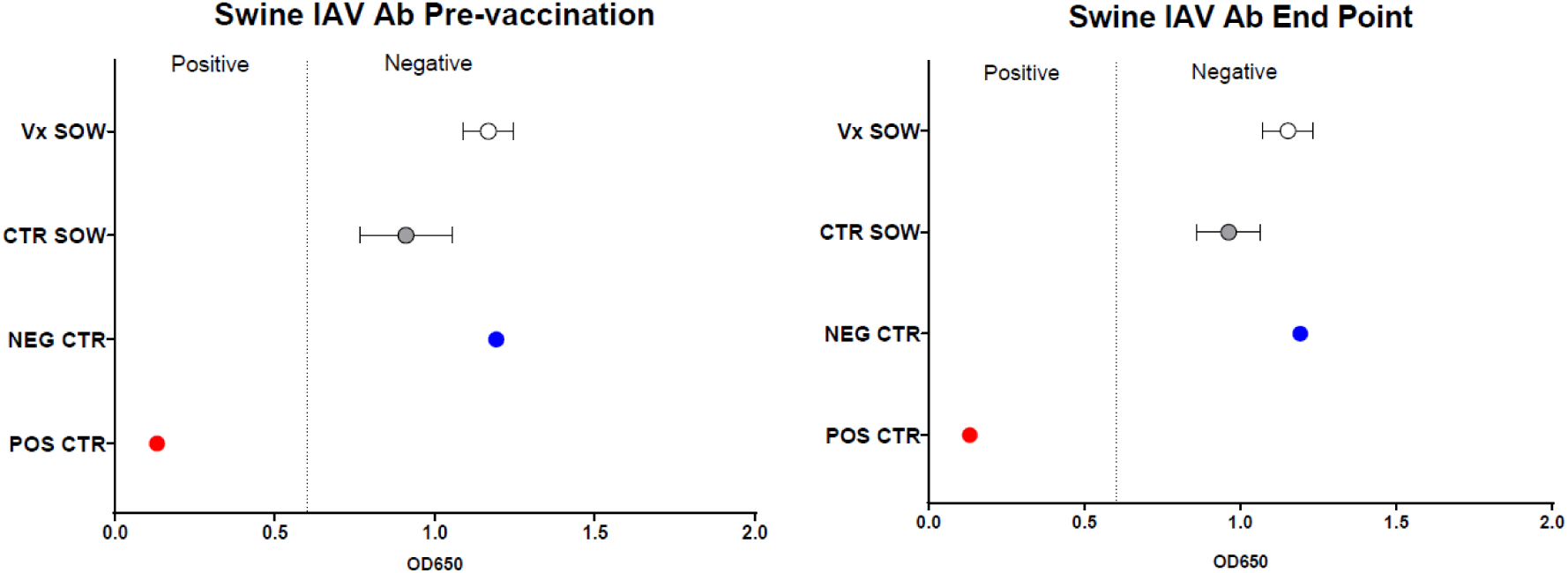
Detection of influenza A virus–specific antibodies. Antibodies were measured using the IDEXX Swine Influenza A Virus (IAV) Antibody ELISA targeting the nucleoprotein (NP), performed according to the manufacturer’s instructions. Samples with a sample-to-negative (S/N) ratio < 0.6 were considered positive.

**Supplementary Figure 2:**
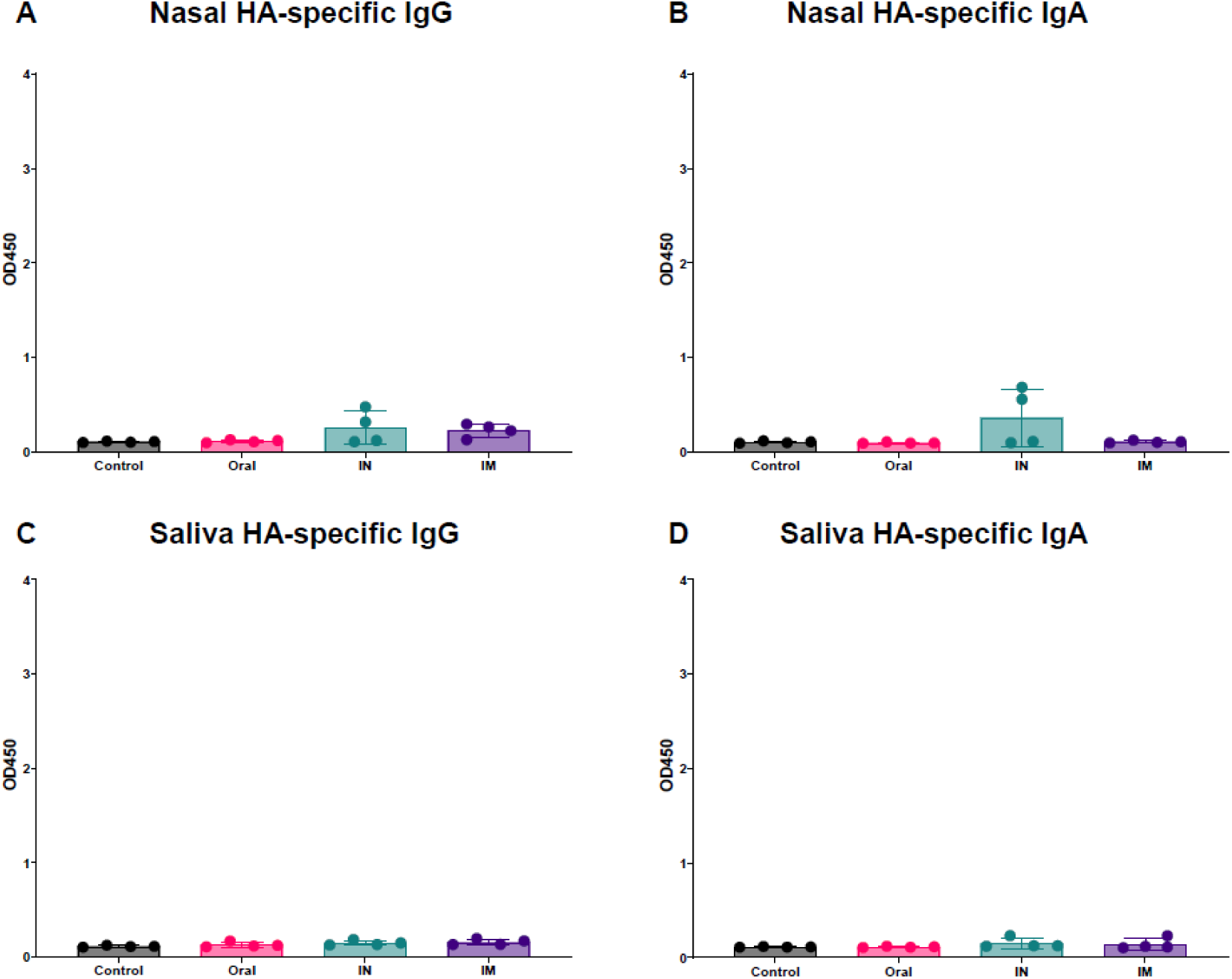
HA-specific IgG and IgA in nasal and oral secretions were not significantly increased by any vaccination route. HA-specific IgG and IgA antibody levels in nasal secretions (A, B) and saliva (C, D) were quantified by ELISA. Data are presented as mean ± SEM. Statistical significance was determined using a two-way ANOVA with Bonferroni’s multiple-comparison correction.

**Supplementary figure 3.**
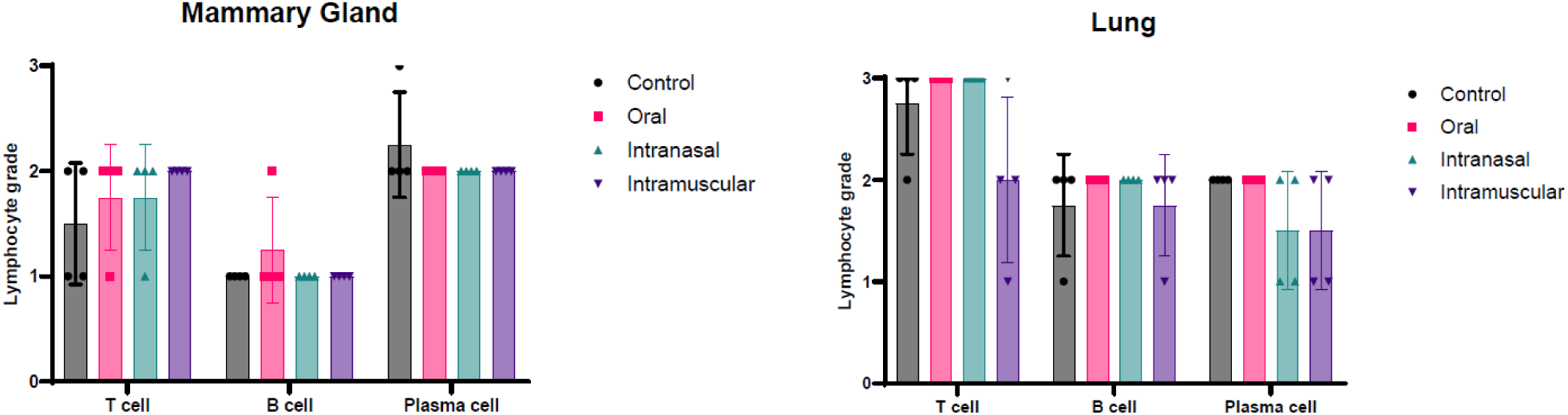
Immunohistochemistry. CD3 and CD20 immunohistochemistry was performed on mammary gland and lung sections. Lymphocyte counts (T, B, or plasma cells) were averaged across 10 high-power fields (hpf) per tissue type. Plasma cells were distinguished from other lymphocytes and enumerated on H&E-stained slides. Grade 0 = No T/B cells or plasma cells present in any hpf, Grade 1 = <10 cells per hpf, Grade 2 = 10-50 cells per hpf, Grade 3 = >50 cells per hpf.

## 2. Author Contributions

SL: Writing – original draft, Writing – review & editing, Validation, Resources, Project administration, Funding acquisition, Formal analysis

JJB: Data curation, Formal analysis, Investigation, Methodology, Validation, Visualization, Writing – review & editing

DL: Investigation, Writing – review & editing

CS: Investigation, Formal analysis, Writing – review & editing AW: Investigation, Formal analysis, Writing – review & editing

DM: Investigation, Formal analysis, Formal analysis, Writing – review & editing

MR: Investigation, Writing – review & editing, Formal analysis TW: Investigation, Writing – review & editing

SC: Investigation, Formal analysis, Data curation, Writing – review & editing DSR: Investigation, Formal analysis, Data curation, Writing – review & editing JBF: Investigation, Writing – review & editing

SNT: Resources, Writing – review & editing

EC: Conceptualization, Data curation, Formal analysis, Funding acquisition, Investigation, Project administration, Resources, Supervision, Validation, Visualization, Writing – original draft, Writing – review & editing

## 4. Acknowledgments

We thank Dr Flowers and the NCSU Swine Educational Unit staff and managers, the NC State College of Veterinary Medicine Laboratory Animal Resources and CPL staff members who assisted in animal husbandry and sampling during the study. We also thank Dr Sitthicharoenchai for helping with animal sampling.

## 5. Data Availability Statement

The raw data supporting the conclusions of this article will be made available by the authors, without undue reservation.

## References

1. Heikkinen, T., H. Silvennoinen, V. Peltola, T. Ziegler, R. Vainionpää, T. Vuorinen, L. Kainulainen, T. Puhakka, T. Jartti, P. Toikka, P. Lehtinen, T. Routi, and T. Juvén. 2004. Burden of influenza in children in the community. J Infect Dis 190: 1369–1373.

2. Izurieta, H. S., W. W. Thompson, P. Kramarz, D. K. Shay, R. L. Davis, F. DeStefano, S. Black, H. Shinefield, and K. Fukuda. 2000. Influenza and the rates of hospitalization for respiratory disease among infants and young children. New Engl J Med 342: 232–239.

3. Lindsay, L., L. A. Jackson, D. A. Savitz, D. J. Weber, G. G. Koch, L. Kong, and H. A. Guess. 2006. Community influenza activity and risk of acute influenza-like illness episodes among healthy unvaccinated pregnant and postpartum women. Am J Epidemiol 163: 838–848.

4. Louie, J. K., M. Acosta, D. J. Jamieson, M. A. Honein, and G. California Pandemic Working. 2010. Severe 2009 H1N1 influenza in pregnant and postpartum women in California. N Engl J Med 362: 27–35.

5. 2012. Vaccines against influenza WHO position paper – November 2012. Wkly Epidemiol Rec 87: 461–476.

6. Grohskopf, L. A., L. Z. Sokolow, K. R. Broder, E. B. Walter, J. S. Bresee, A. M. Fry, and D. B. Jernigan. 2017. Prevention and Control of Seasonal Influenza with Vaccines: Recommendations of the Advisory Committee on Immunization Practices - United States, 2017-18 Influenza Season. MMWR Recomm Rep 66: 1–20.

7. Groothuis, J. R., M. J. Levin, G. P. Rabalais, G. Meiklejohn, and B. A. Lauer. 1991. Immunization of high-risk infants younger than 18 months of age with split-product influenza vaccine. Pediatrics 87: 823–828.

8. Zangwill, K. M., and R. B. Belshe. 2004. Safety and efficacy of trivalent inactivated influenza vaccine in young children: a summary for the new era of routine vaccination. Pediatr Infect Dis J 23: 189–197.

9. Shang, M., L. Blanton, L. Brammer, S. J. Olsen, and A. M. Fry. 2018. Influenza-Associated Pediatric Deaths in the United States, 2010-2016. Pediatrics 141.

10. Hurley, W. L., and P. K. Theil. 2011. Perspectives on immunoglobulins in colostrum and milk. Nutrients 3: 442–474.

11. Blanchard-Rohner, G., S. Meier, M. Bel, C. Combescure, V. Othenin-Girard, R. A. Swali, B. Martinez de Tejada, and C. A. Siegrist. 2013. Influenza vaccination given at least 2 weeks before delivery to pregnant women facilitates transmission of seroprotective influenza-specific antibodies to the newborn. Pediatr Infect Dis J 32: 1374–1380.

12. Madhi, S. A., C. L. Cutland, L. Kuwanda, A. Weinberg, A. Hugo, S. Jones, P. V. Adrian, N. van Niekerk, F. Treurnicht, J. R. Ortiz, M. Venter, A. Violari, K. M. Neuzil, E. A. Simões, K. P. Klugman, and M. C. Nunes. 2014. Influenza vaccination of pregnant women and protection of their infants. N Engl J Med 371: 918–931.

13. Zuccotti, G., L. Pogliani, E. Pariani, A. Amendola, and A. Zanetti. 2010. Transplacental antibody transfer following maternal immunization with a pandemic 2009 influenza A(H1N1) MF59-adjuvanted vaccine. Jama 304: 2360–2361.

14. Schlaudecker, E. P., M. C. Steinhoff, S. B. Omer, M. M. McNeal, E. Roy, S. E. Arifeen, C. N. Dodd, R. Raqib, R. F. Breiman, and K. Zaman. 2013. IgA and neutralizing antibodies to influenza a virus in human milk: a randomized trial of antenatal influenza immunization. PLoS One 8: e70867.

15. Morteau, O., C. Gerard, B. Lu, S. Ghiran, M. Rits, Y. Fujiwara, Y. Law, K. Distelhorst, E. M. Nielsen, E. D. Hill, R. Kwan, N. H. Lazarus, E. C. Butcher, and E. Wilson. 2008. An indispensable role for the chemokine receptor CCR10 in IgA antibody-secreting cell accumulation. J Immunol 181: 6309–6315.

16. Wilson, E., and E. C. Butcher. 2004. CCL28 controls immunoglobulin (Ig)A plasma cell accumulation in the lactating mammary gland and IgA antibody transfer to the neonate. J Exp Med 200: 805–809.

17. Cox, R. J., K. A. Brokstad, and P. Ogra. 2004. Influenza virus: immunity and vaccination strategies. Comparison of the immune response to inactivated and live, attenuated influenza vaccines. Scand J Immunol 59: 1–15.

18. Neutra, M. R., and P. A. Kozlowski. 2006. Mucosal vaccines: the promise and the challenge. Nat Rev Immunol 6: 148–158.

19. Meurens, F., A. Summerfield, H. Nauwynck, L. Saif, and V. Gerdts. 2012. The pig: a model for human infectious diseases. Trends Microbiol 20: 50–57.

20. Lunney, J. K., A. Van Goor, K. E. Walker, T. Hailstock, J. Franklin, and C. Dai. 2021. Importance of the pig as a human biomedical model. Sci Transl Med 13: eabd5758.

21. Chattha, K. S., J. A. Roth, and L. J. Saif. 2015. Strategies for design and application of enteric viral vaccines. Annu Rev Anim Biosci 3: 375–395.

22. Liebowitz, D., K. Gottlieb, N. S. Kolhatkar, S. J. Garg, J. M. Asher, J. Nazareno, K. Kim, D. R. McIlwain, and S. N. Tucker. 2020. Efficacy, immunogenicity, and safety of an oral influenza vaccine: a placebo-controlled and active-controlled phase 2 human challenge study. Lancet Infect Dis 20: 435–444.

23. Braun, M. R., L. Q. Nguyen, B. A. Flitter, N. J. Bennett, D. C. M. Hailey, C. A. Lester, E. D. Neuhaus, K. Marx, N. P. D’Amato, K. Tam, M. F. Pasetti, S. N. Tucker, and J. F. Cummings. 2026. Transfer of breast milk IgA to infants after oral bivalent norovirus vaccination of post-partum women. NPJ Vaccines 11: 44.

24. De Rensis, F., R. Saleri, P. Tummaruk, M. Techakumphu, and R. N. Kirkwood. 2012. Prostaglandin F2α and control of reproduction in female swine: A review. Theriogenology 77: 1–11.

25. Rahe, M. C., J. J. Byrne, D. Meritet, E. Gruber, S. N. Langel, and E. Crisci. 2025. Case Report: The effect of asplenia on the response to influenza vaccination and passive transfer of immunity in an adult female pig. Frontiers in Immunology Volume 16 - 2025.

26. Lei, C., J. Yang, J. Hu, and X. Sun. 2021. On the Calculation of TCID(50) for Quantitation of Virus Infectivity. Virol Sin 36: 141–144.

27. Kitikoon, P., P. C. Gauger, and A. L. Vincent. 2014. Hemagglutinin inhibition assay with swine sera. Methods Mol Biol 1161: 295–301.

28. Crisci, E., A. R. Kick, L. M. Cortes, J. J. Byrne, A. F. Amaral, K. Love, H. Tong, J. Zhang, P. C. Gauger, J. S. Pittman, and T. Kaser. 2025. Challenges and Lessons Learned from a Field Trial on the Understanding of the Porcine Respiratory Disease Complex. Vaccines (Basel) 13.

29. Kollmann, T. R., A. Marchant, and S. S. Way. 2020. Vaccination strategies to enhance immunity in neonates. Science 368: 612–615.

30. Langel, S. N., M. Blasi, and S. R. Permar. 2022. Maternal immune protection against infectious diseases. Cell Host Microbe 30: 660–674.

31. Afkhami, S., Y. Yao, and Z. Xing. 2016. Methods and clinical development of adenovirus-vectored vaccines against mucosal pathogens. Mol Ther Methods Clin Dev 3: 16030.

32. Sakurai, F., M. Tachibana, and H. Mizuguchi. 2022. Adenovirus vector-based vaccine for infectious diseases. Drug Metab Pharmacokinet 42: 100432.

33. Shao, H. Y., Y. C. Chen, N. H. Chung, Y. J. Lu, C. K. Chang, S. L. Yu, C. C. Liu, and Y. H. Chow. 2018. Maternal immunization with a recombinant adenovirus-expressing fusion protein protects neonatal cotton rats from respiratory syncytia virus infection by transferring antibodies via breast milk and placenta. Virology 521: 181–189.

34. Song, X., Q. Zhou, J. Zhang, T. Chen, G. Deng, H. Yue, C. Tang, X. Wu, J. Yu, and B. Zhang. 2024. Immunogenicity and protective efficacy of recombinant adenovirus expressing a novel genotype G2b PEDV spike protein in protecting newborn piglets against PEDV. Microbiol Spectr 12: e0240323.

35. Choi, K. Y., Y. Qin, N. El-Hamdi, and A. McGregor. 2025. Complete cross strain protection against congenital cytomegalovirus infection requires a vaccine encoding key antibody (gB) and T-cell (immediate early 1 protein) viral antigens. Front Immunol 16: 1649656.

36. Kim, E., G. Erdos, S. Huang, T. Kenniston, L. D. Falo, Jr., and A. Gambotto. 2016. Preventative Vaccines for Zika Virus Outbreak: Preliminary Evaluation. EBioMedicine 13: 315–320.

37. Jia, S., Y. Liu, Q. He, H. Pan, Z. Liang, J. Zhou, Y. Pan, S. Liu, J. Wu, K. Yang, X. Zhang, Y. Zhao, S. Li, L. Zhang, L. Chen, A. Yao, M. Lu, F. Zhu, Q. Mao, and J. Li. 2025. Effectiveness of a booster dose of aerosolized or intramuscular adenovirus type 5 vectored COVID-19 vaccine in adults: a multicenter, partially randomized, platform trial in China. Nat Commun 16: 2969.

38. Tang, R., H. Zheng, B. S. Wang, J. B. Gou, X. L. Guo, X. Q. Chen, Y. Chen, S. P. Wu, J. Zhong, H. X. Pan, J. H. Zhu, X. Y. Xu, F. J. Shi, Z. P. Li, J. X. Liu, X. Y. Zhang, L. B. Cui, Z. Z. Song, L. H. Hou, F. C. Zhu, J. X. Li, and C.-S. G. CanSino. 2023. Safety and immunogenicity of aerosolised Ad5-nCoV, intramuscular Ad5-nCoV, or inactivated COVID-19 vaccine CoronaVac given as the second booster following three doses of CoronaVac: a multicentre, open-label, phase 4, randomised trial. Lancet Respir Med 11: 613–623.

39. Torres, J. M., C. Alonso, A. Ortega, S. Mittal, F. Graham, and L. Enjuanes. 1996. Tropism of human adenovirus type 5-based vectors in swine and their ability to protect against transmissible gastroenteritis coronavirus. J Virol 70: 3770–3780.

40. Corthésy, B. 2013. Multi-faceted functions of secretory IgA at mucosal surfaces. Front Immunol 4: 185.

41. Rajao, D. S., G. C. Zanella, M. Wymore Brand, S. Khan, M. E. Miller, L. M. Ferreri, C. J. Caceres, S. Cadernas-Garcia, C. K. Souza, T. K. Anderson, P. C. Gauger, A. L. Vincent Baker, and D. R. Perez. 2023. Live attenuated influenza A virus vaccine expressing an IgA-inducing protein protects pigs against replication and transmission. Frontiers in Virology Volume 3–2023.

42. Baker, P. H., L. S. Stafford, and S. N. Langel. 2025. Advancing protective effects of maternal antibodies in neonates through animal models. J Immunol 214: 2523–2534.

43. Bourne, F. J., and J. Curtis. 1973. The transfer of immunoglobins IgG, IgA and IgM from serum to colostrum and milk in the sow. Immunology 24: 157–162.

44. Verschoor, C. P., P. Singh, M. L. Russell, D. M. Bowdish, A. Brewer, L. Cyr, B. J. Ward, and M. Loeb. 2015. Microneutralization assay titres correlate with protection against seasonal influenza H1N1 and H3N2 in children. PLoS One 10: e0131531.

45. Stephenson, I., R. G. Das, J. M. Wood, and J. M. Katz. 2007. Comparison of neutralising antibody assays for detection of antibody to influenza A/H3N2 viruses: an international collaborative study. Vaccine 25: 4056–4063.

